# Enhanced representation of natural sound sequences in the ventral auditory midbrain

**DOI:** 10.1101/846485

**Authors:** Eugenia González-Palomares, Luciana López-Jury, Francisco García-Rosales, Julio C. Hechavarria

## Abstract

**Summary:** The auditory midbrain (inferior colliculus, IC) plays an important role in sound processing, acting as hub for acoustic information extraction and for the implementation of fast audio-motor behaviors. IC neurons are topographically organized according to their sound frequency preference: dorsal IC regions encode low frequencies while ventral areas respond best to high frequencies, a type of sensory map defined as tonotopy. Tonotopic maps have been studied extensively using artificial stimuli (pure tones) but our knowledge of how these maps represent information about sequences of natural, spectro-temporally rich sounds is sparse. We studied this question by conducting simultaneous extracellular recordings across IC depths in awake bats (*Carollia perspicillata*) that listened to sequences of natural communication and echolocation sounds. The hypothesis was that information about these two types of sound streams is represented at different IC depths since they exhibit large differences in spectral composition, i.e. echolocation covers the high frequency portion of the bat soundscape (> 45 kHz), while communication sounds are broadband and carry most power at low frequencies (20-25 kHz). Our results showed that mutual information between neuronal responses and acoustic stimuli, as well as response redundancy in pairs of neurons recorded simultaneously, increase exponentially with IC depth. The latter occurs regardless of the sound type presented to the bats (echolocation or communication). Taken together, our results indicate the existence of mutual information and redundancy maps at the midbrain level whose response cannot be predicted based on the frequency composition of natural sounds and classic neuronal tuning curves.

## Introduction

Animals depend greatly on acoustic signals to interact with the environment and other life beings. Encoding acoustic information in the auditory system is a fundamental step leading to the production of behavioral responses in everyday scenarios (Brudzynski, 2013; Jiang et al., 2017; Liévin-Bazin et al., 2018; Ryan et al., 1985). The latter could determine the animals’ well-being and their capacity to adapt to environmental pressures.

The main aim of this article is to study the representation of natural sounds in the auditory midbrain (inferior colliculus, IC). The IC is an important integration hub in the auditory pathway that has been linked to the production of fast audio-motor behaviors instrumental for animal survival (Casseday and Covey, 1996; Covey et al., 1987; Malmierca, 2004). This structure is also a target area for auditory prostheses that benefit deaf patients who cannot sufficiently profit from cochlear implants (Colletti et al., 2007; Lim et al., 2007, 2008). Though the IC has been studied extensively at the anatomical and functional levels (Casseday et al., 2002; Malmierca, 2004), our knowledge of how neurons within this structure represent natural sound streams is still sparse.

We relied on bats as experimental animal model to assess how natural sound sequences are represented simultaneously across IC depths. Bats represent an excellent animal model for auditory experiments because of their rich soundscape, which includes echolocation (sound-based navigation) and multiple types of communication sounds (Schnitzler et al., 2003; Wilkinson and Boughman, 1998). The latter are used to maintain hierarchies in the colonies, to communicate with infants and to alert other individuals about potentially dangerous/uncomfortable situations (Balcombe and McCracken, 1992; Gadziola et al., 2012; Knörnschild et al., 2013).

The auditory system of bats has been heavily studied in the last decades but, at present, no consensus exists as to whether communication and echolocation sounds can be represented by the same neurons (Kössl et al., 2015). In the bat species *Carollia perspicillata* (the species of choice for this study), there is a clear dissociation in the frequency domain between communication and biosonar sounds used for orientation. The former cover the low frequency portion of the bat soundscape, with the power of individual syllables peaking at frequencies close to 20 kHz, while the latter carry most energy in the high frequency band between 45-100 kHz (Fig. 1; Brinkløv et al., 2011; Hechavarría et al., 2016).

**Figure 1.**
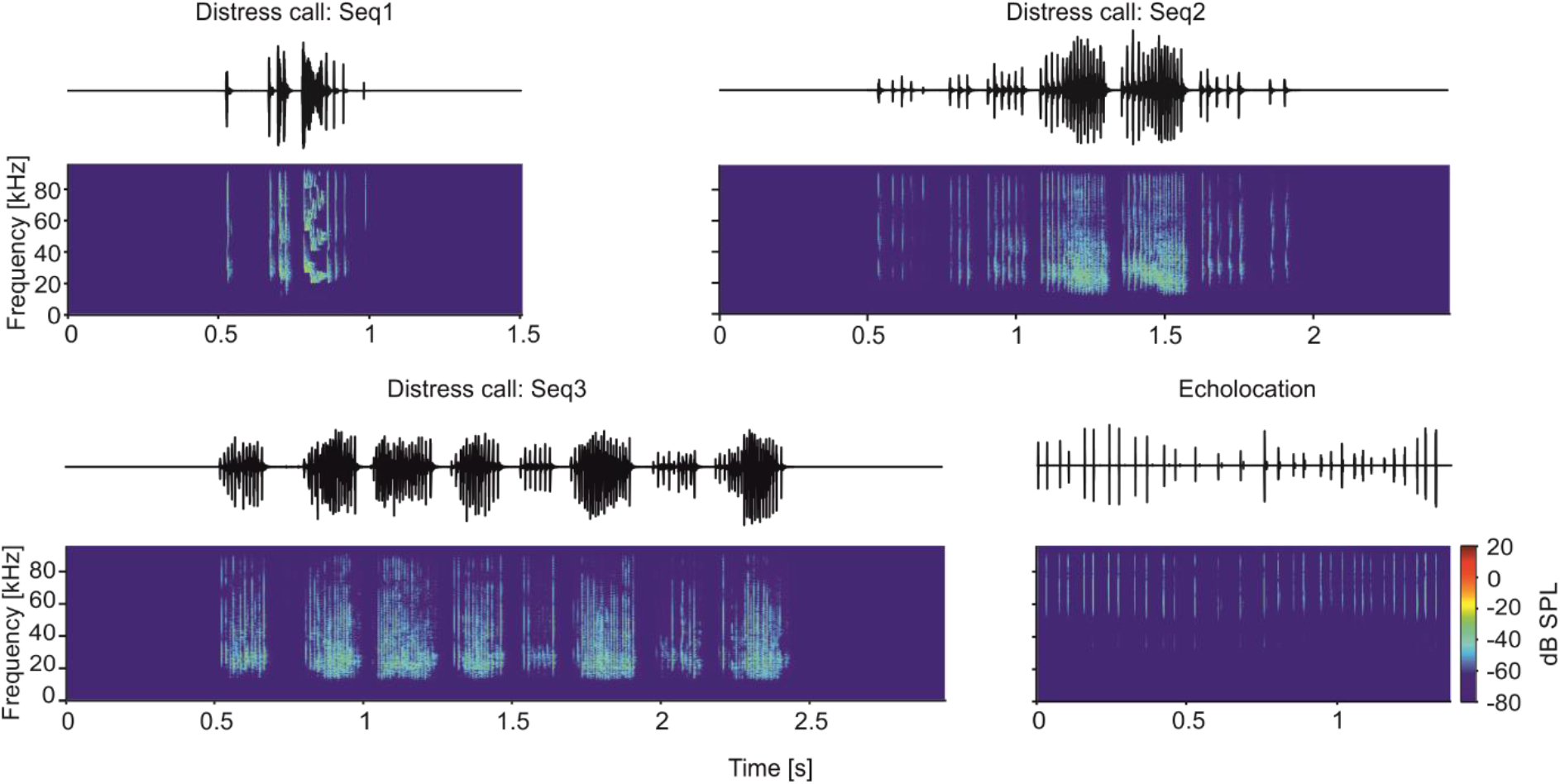
Oscillograms and spectrograms of the natural calls used as stimuli, three distress calls (Seq1, Seq2 and Seq3) and one biosonar call (Echolocation).

The bat IC follows the general mammalian plan with a dorso-ventral tonotopic arrangement in which neurons located close to the brain surface process low frequencies, and neurons located in ventral IC layers process high frequencies (Friauf, 1992; Grinnell, 1963; Jen and Chen, 1998; Malmierca et al., 2008). There is one peculiarity in *C. perspicillata*’s IC: although ventral neurons are responsive to high frequencies they respond as well to low frequency sounds (Beetz et al., 2017). In other words, neurons located in the ventral IC of this bat species display multipeaked frequency tuning curves and all neurons throughout the IC’s tonotopy respond (at least to some extent) to sounds whose carrier frequency lies in the 20-30 kHz range. It has been speculated that multipeaked frequency tuning could allow neurons to respond to both, echolocation and communication sounds (Kanwal, Jagmeet and Rauschecker, 2007; Kössl et al., 2015). We reasoned that distress utterances produced by *C. perspicillata* could drive activity throughout the entire IC, since both dorsal and ventral neurons are responsive to frequencies ~ 20 kHz, corresponding to the peak frequencies of distress vocalizations (Hechavarría et al., 2016a).

Using laminar probes we performed simultaneous recordings from dorsal and ventral IC areas in awake *C. perspicillata*. Our hypothesis was that, in response to echolocation sequences, the information provided by collicular neurons should be highest in ventral IC regions since echolocation sounds are mostly high frequency. On the other hand, information about communication sequences could be either highest in dorsal IC or equally distributed throughout the entire structure, due to the presence of multipeaked frequency tuning curves in ventral areas. The data revealed that ventral IC regions are more informative than dorsal regions not only about echolocation sounds, but also about communication, which was surprising given our original hypothesis. The ventral IC also contains the highest degree of response redundancy in pairs of neurons recorded simultaneously. This redundancy is tightly linked to signal correlations in the neurons recorded. Overall, the data presented in this article provides evidence for topographical representations of acoustic information and redundancy in the mammalian midbrain in naturalistic contexts.

## Results

The activity of 864 units (one per electrode and penetration) was recorded from the central nucleus of the inferior colliculus (IC) in awake bats (species *C. perspicillata*) in response to pure tones and to four natural sound sequences (Fig. 1, for sequence parameters see Table 1). Natural sequences consisted of three distress calls (*Seq1-3*) and one echolocation sequence. Distress calls are a type of communication sound used to advertise danger/discomfort to others (Eckenweber and Knörnschild, 2016; Hechavarría et al., 2016a; Russ et al., 2004, 1998). The three distress sequences were chosen because they constitute typical examples of bats’ alarm utterances (Hechavarría et al., 2016a). Only one biosonar sequence was used, as echolocation is a stereotyped behavior that involves fixed action patterns as bats approach a target (Neuweiler, 1990, 2003; Thies et al., 1998). The echolocation sequence was recorded in a pendulum paradigm in which a bat was swung towards a reflective wall (Beetz et al., 2016a). Distress and echolocation sequences have been used in previous studies characterizing the bat auditory system (Beetz et al., 2016a, 2017; García-Rosales et al., 2018; Hechavarría et al., 2016b; Macías et al., 2018; Martin et al., 2017; Wohlgemuth and Moss, 2016). To restrict the study to units that responded to the calls, we considered only those units that carried at least 1 bit/s of information (Kayser et al., 2009) in response to at least one of the sequences studied (864 units out of 976).

**Table 1.** Basic temporal properties of the natural distress sequences used as stimuli. The sound intensity was obtained considering each syllable individually per each sequence.

| Seq | # of bouts | # of syllables | Sequence length [s] | Avg. length of bouts [ms] | Avg. length syllables [ms] | Avg. inter-syllable interval [ms] | Intensity (mean ± STD) [db SPL] |
| --- | --- | --- | --- | --- | --- | --- | --- |
| 1 | 1 | 9 | 0.51 | 510 | 69.2 | 141.4 | 90.2 ± 5.06 |
| 2 | 7 | 54 | 1.46 | 140 | 4.8 | 26.4 | 90.6 ± 2.45 |
| 3 | 8 | 122 | 1.96 | 176.6 | 2.9 | 15.7 | 90.2 ± 1.93 |
| Echo | / | 31 | 1.38 | / | 0.9 | 44.1 | 88.4 ± 5.0 |

### General properties of *C. perspicillata*’s auditory midbrain

Iso-level frequency tuning curves (FTCs) were analyzed to confirm the tonotopy along the dorso-ventral axis of IC’s central nucleus (Beetz et al., 2017; Friauf, 1992; Grinnell, 1963; Jen and Chen, 1998; Malmierca et al., 2008). The inferior colliculus is functionally organized in iso-frequency layers, with each layer being sensitive to a narrow range of frequencies, from low to high frequencies in the dorso-ventral axis (Friauf, 1992; Malmierca et al., 2008). We analyzed the response to pure tones (frequencies from 10 to 90 kHz, steps of 5 kHz, 60 dB SPL) in terms of number of spikes to create iso-level FTCs. Fig. 2A shows the results of an example recording and the iso-level FTCs obtained in all 16 channels simultaneously. Two peaks of high activity are evident, especially in deep IC areas. The high-frequency peak shifts to higher frequencies as the depth of the channels increases which demonstrates the tonotopy of the inferior colliculus (Fig. 2B, population data). The low-frequency peak (10-30 kHz) occurs throughout all depths studied. Note that double-peaked FTCs have been reported before in the bats’ IC (Beetz et al., 2017; Casseday and Covey, 1992; Holmstrom et al., 2007; Mittmann and Wenstrup, 1995), as well as in the AC of bats and other species (Fitzpatrick et al., 1993; Hagemann et al., 2011; Sutter and Schreiner, 1991) and frontal auditory areas (López-Jury et al., 2019).

**Figure 2.**
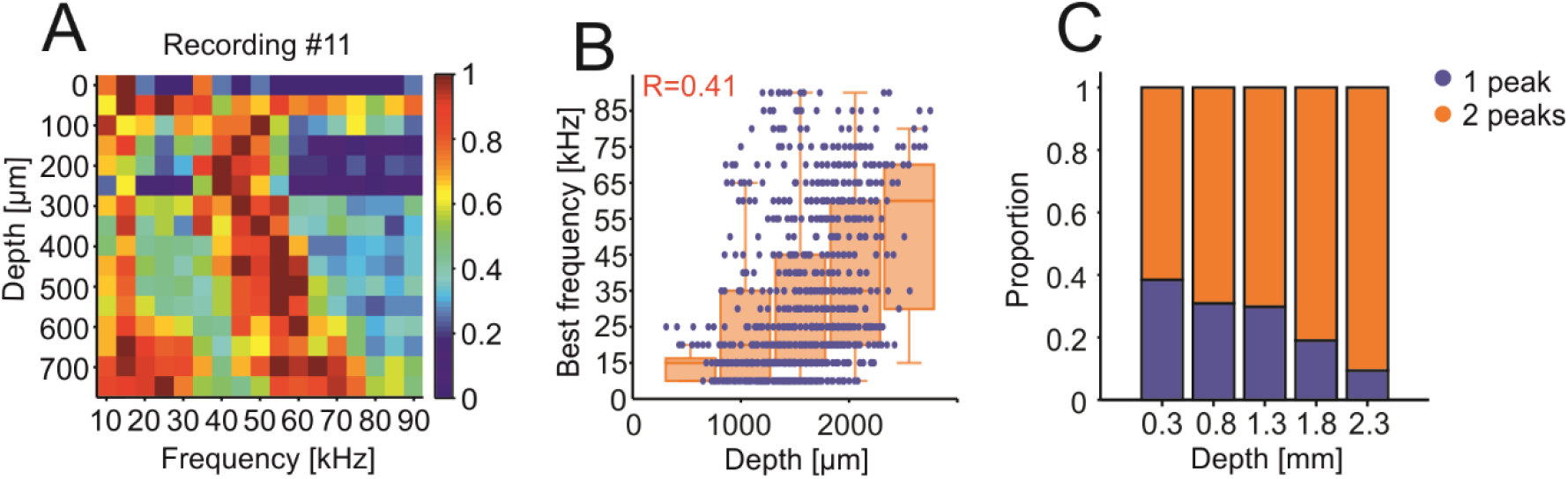
Tonotopy in the inferior colliculus. A) Normalized (for each channel) number of spikes of the recording #11 plotted against the channel depth (relative to the most dorsal channel). B) Scatter plot of the depth and best frequency of each unit. Note the general increase in best frequency as the depth increases. In red, correlation coefficient (R) of the exponential curve fitted into the data with the bisquare robust method and equation: *f*(*x*) = 1.2*e*^0.0009*x*^. Superimposed are the mean + SEM of the best frequency per discretized depth (each bar spans 0.5 mm starting at 0.3 mm depth). C) Proportion of single-peak and double-peak iso-level FTCs. Single-peak units had a peak only in the range 10-45 kHz or 50-90 kHz. Double-peak units had peaks in both frequency ranges (peak defined as > 60% of the maximum spike-count value). Note the proportional increase of double-peak units in more ventral regions.

As expected from the known tonotopy of the IC, there was a positive correlation between neuronal best frequency (i.e., frequency that triggers the largest number of spikes) and the depth of the channel that recorded the units (Fig. 2B). In addition, as IC depth increased, so did the probability of finding units with complex FTCs having more than one peak (multi-peaked FTCs, Fig. 2A and C). At the population level, there was an overrepresentation of low best frequencies (10-30 kHz), likely influenced by the fact that neurons tuned to high frequencies were also responsive to low frequency tones (Fig. S1A). Note that the range from 10-30 kHz corresponds to the peak frequencies in distress calls (Figs. 1 and S1B).

Based on the results obtained with pure tones, one could speculate that distress sounds (with peak frequencies at 20-30 kHz) should be best represented in dorsal IC layers or throughout the extent of the IC. On the other hand, echolocation sounds should drive strongest spiking in deep IC regions responsive to high frequencies.

### Ventral units in the inferior colliculus are better trackers of natural auditory streams

The main aim of this study was to assess whether there is a difference in information representation in response to natural sounds across IC depths. As stimuli, we used natural distress sequences, which carry most power in low frequencies (20-30 kHz), and an echolocation sequence with high power in frequencies ranging from 60-90 kHz (see stimuli in Fig. 1 and stimuli spectra in supplementary Figure S1B).

A qualitative check of the neural responses to the sequences already revealed that dorsal units were worse in representing natural sound streams than ventral units, regardless of the type of stimulus presented (distress or echolocation). Ventral units appear more precise and reliable across trials in their responses to both distress and echolocation sequences (see example dorsal and ventral units in Fig. 3B and C, respectively).

**Figure 3.**
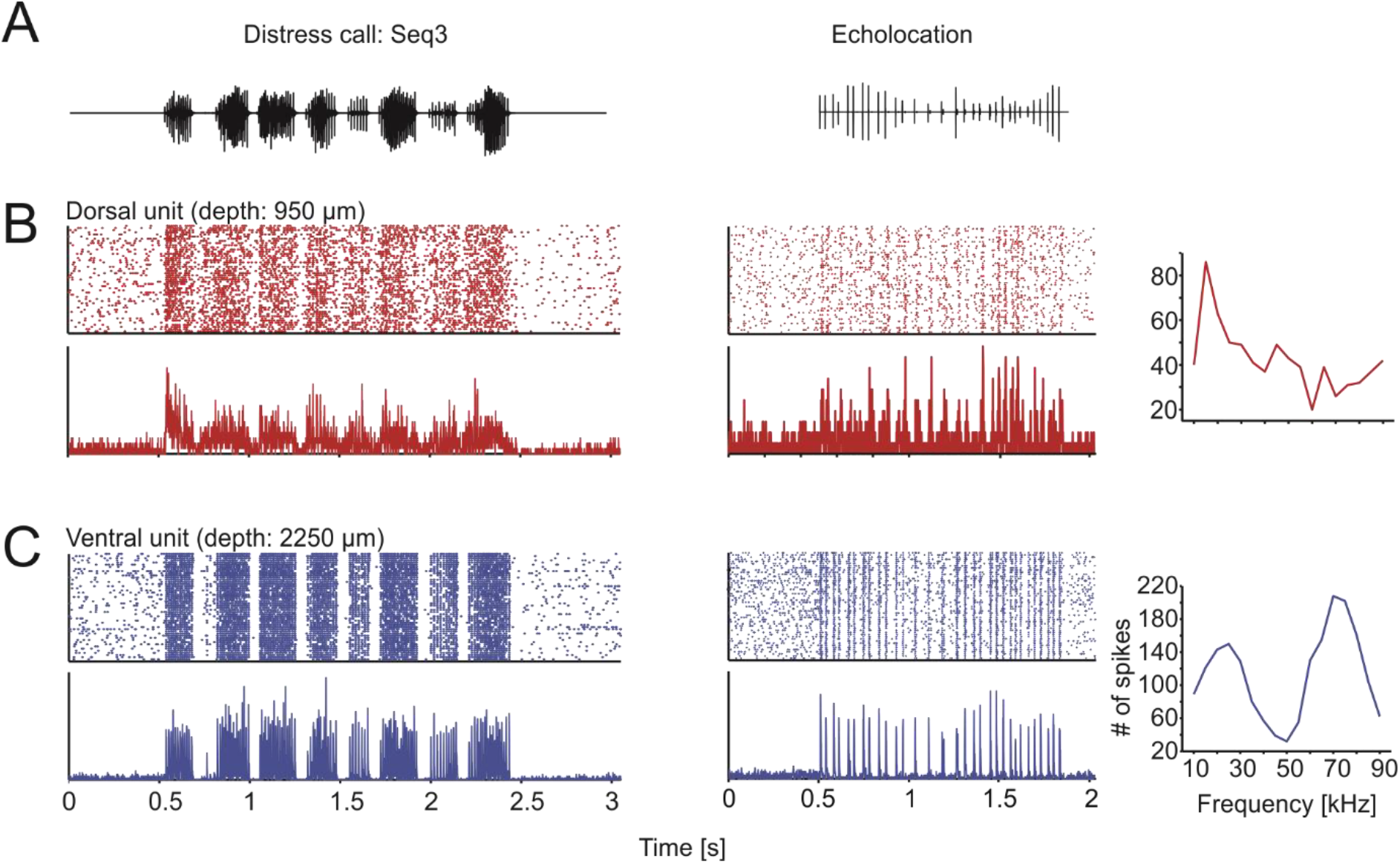
Ventral units represent more accurately the stimuli. A) Oscillograms of the natural calls used as stimuli. B) Raster plots (50 trials in total; top left) and peristimulus time histogram (PSTH; 1 ms precision; bottom) of one exemplary dorsal unit and one ventral unit in response to the sequences shown in A. Note the higher precision and reliability in the ventral unit. On the right are the frequency tuning curves of each unit. While the dorsal unit has a clear peak in the low frequencies, the ventral unit shows a double-peaked curve, one low- and the other high-frequency peak.

Differences between dorsal and ventral units regarding the information they provide about natural sound streams were quantified by means of Shannon’s mutual information. Mutual information calculations revealed that the information (*I*_*rate*_) provided by the units increased exponentially with IC depth, regardless of the type of sequence (i.e. distress or echolocation) used as stimulus (Fig. 4). In other words, observing the neuronal response of ventral units reduces more the uncertainty about the stimulus than the response from dorsal units, and this trend was independent of the sounds heard. As shown in Figure 4A, *I*_*rate*_ and IC depth had an exponential, relation, with increasing *I*_*rate*_ with IC depth. In order to statistically compare the *I*_*rate*_ at different depths, we classified the *I*_*rate*_ estimates into five depth groups, and corroborated that the ventral units carry more information than dorsal units (Fig. 4B; FDR-corrected Wilcoxon rank-sum tests, *p* < 0.05) across all sound sequences tested.

**Figure 4.**
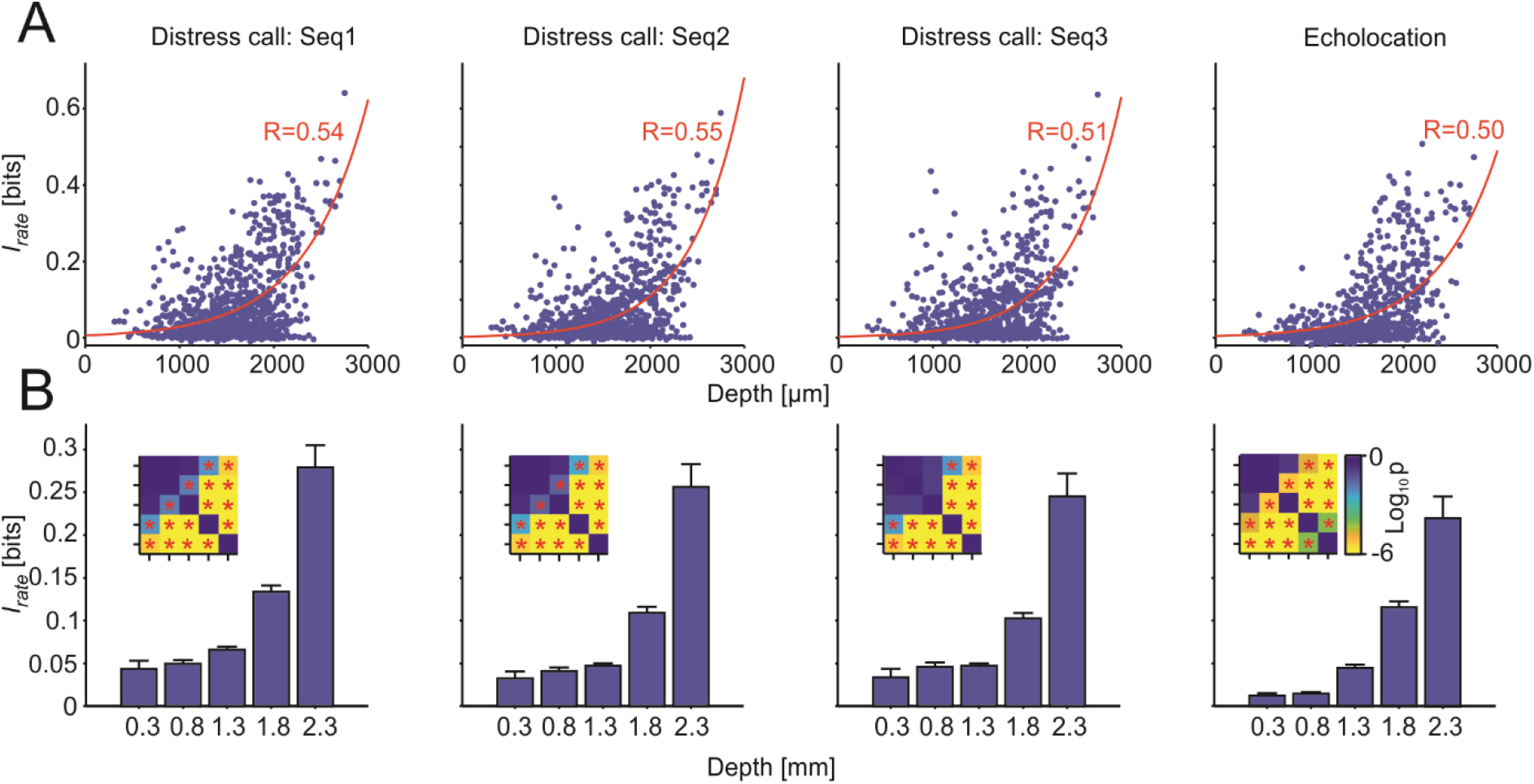
Ventral units carry the most information regardless of the stimuli. A) Scatter plots of the information in the rate code plotted against the depth of the recorded unit for all sequences. In red is the exponential curve fitted to the data with the corresponding correlation coefficient value. B) Mean * SEM of the information as in 4A with discretized depths for all sequences. Each bar represents the information of the units comprised in 0.5 mm depth distance starting at the values stated in the labels. Statistical comparisons were performed by the FDR-corrected Wilcoxon rank-sum tests. The insets show p-value matrices of all the statistical comparisons in a logarithmic scale. * *p*_*corr*_ < 0.05.

Finding an exponential relation between IC depth and mutual information was an unexpected result considering the tonotopic characteristics of the IC and the disparate spectral structure of the stimuli tested (echolocation vs. distress). Our results suggest the presence of a topographical representation of mutual information throughout the IC, presumably linked to the large complexity of receptive fields in ventral IC units (see Discussion).

### Joint information in groups of neurons enables better representation of acoustic stimuli

The information provided by units recorded simultaneously was also quantified by means of joint information calculations (*I*_*joint*_, calculated in a total of 5440 pairs). *I*_*joint*_ measures the information considering responses in pairs of units as their combined activity, considering the identity of individual responses (i.e. which unit fired which spikes, see Methods). Note that a unit can be considered multiple times to form pairs with other units, since we recorded simultaneously from 16 IC positions.

The *I*_*rate*_ estimates of the individual units that composed the pairs were compared to their *I*_*joint*_ (Fig. 5A). This comparison tests whether more information is provided by responses of pairs of units than by each unit separately. The results showed that *I*_*joint*_ was significantly higher than *I*_*rate*_ regardless of the stimulation sequence considered (FDR-corrected Wilcoxon signed-rank tests, p < 0.05). Thus, in the IC, the response of two simultaneously-recorded units provides more information about the stimulus than the response of a single unit. Besides pairs, the information of the spike rate was also calculated for larger groups of units (Figure S3): triplets (n = 16703), quadruplets (n = 31112) and quintuplets (n = 33042). As expected, the information increased with the number of units considered (FDR-corrected Wilcoxon signed-rank tests, p < 0.05), i.e. with higher number of units used to calculate the mutual information, the more uncertainty of the stimulus was reduced.

**Figure 5.**
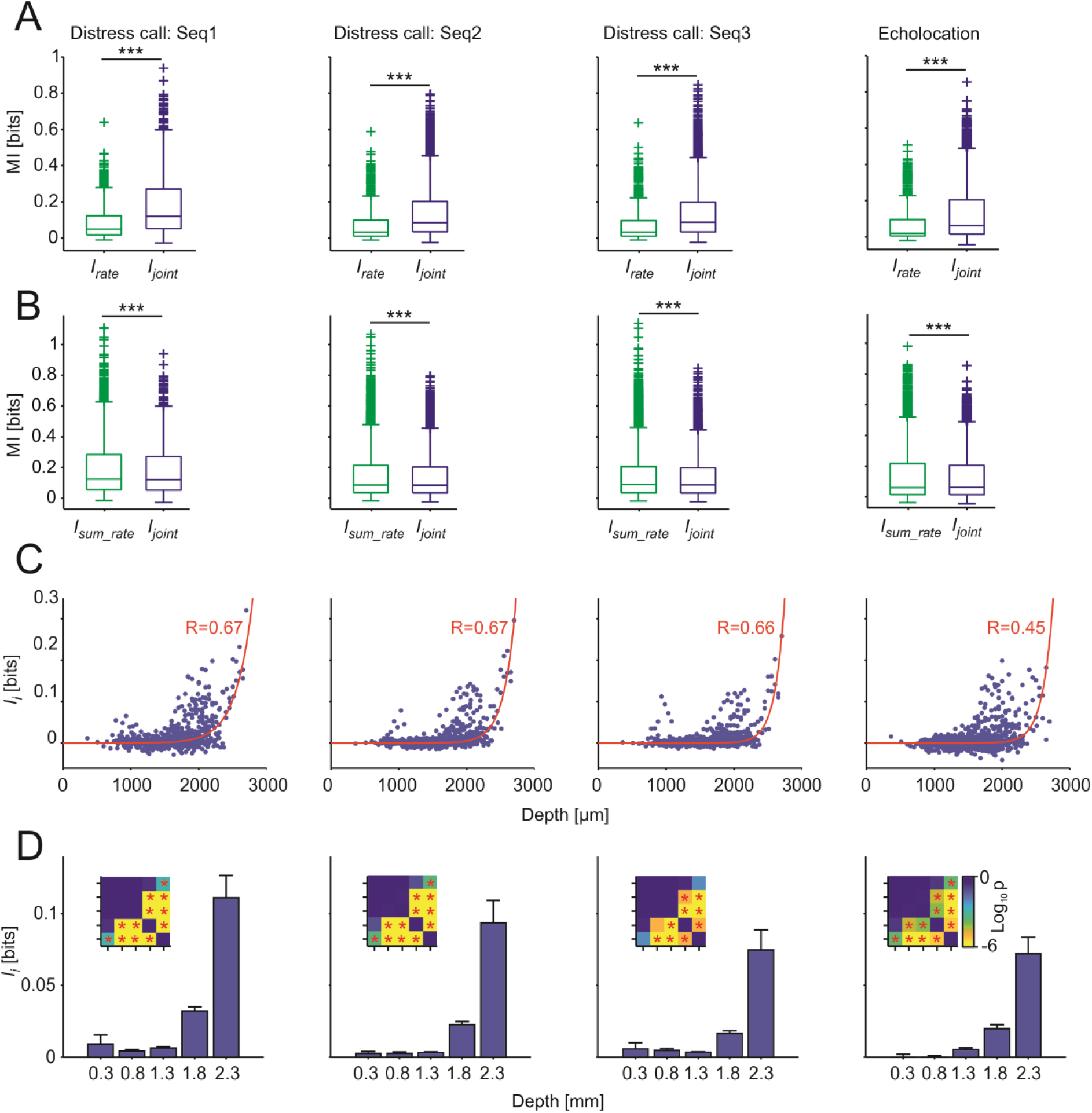
Redundancy increases with depth in simultaneously-recorded units. A) Comparison of *I*_*rate*_ vs *I*_*joint*_ for all units and pairs per sequence. Statistical comparisons performed by FDR-corrected Wilcoxon signed-rank tests. *** *p*_*corr*_ < 0.001. B) Comparison of *I*_*joint*_ vs the sum of information of the rate codes of each of the units that form the pairs (*I*_*sum*_) per sequence. Statistical comparisons performed by FDR-corrected Wilcoxon signed-rank tests. *** *p*_*corr*_ < 0.001. C) Scatter plots of the *I*_*i*_ estimates (*I*_*sum*_ − *I*_*joint*_; *I*_*sum*_ = *I*_*rate(a)*_ + *I*_*rate(b)*_) plotted against the intermediate depth of the pairs (only those pairs with units distanced by 100 μm were considered). In red is the exponential curve fitted to the data with the corresponding correlation coefficient value. D) Mean + SEM of the *I*_*i*_ as in C with discretized depths for all sequences. Each bar represents the *I*_*i*_ of the pairs comprised in 0.5 mm depth distance starting at the values stated in the labels. Statistical comparisons were performed by the FDR-corrected Wilcoxon rank-sum tests. The insets show p-value matrices of all the statistical comparisons in a logarithmic scale. * *p*_*corr*_ < 0.05.

### An information redundancy map exists in the auditory midbrain

To estimate if pairs of units carried redundant information, *I*_*joint*_ was statistically compared to the linear sum of *I*_*rate*_ of the units that formed the pairs (*I*_*sum*_). *I*_*sum*_ was significantly higher than *I*_*joint*_ in all the sequences considered (Fig. 5B; FDR-corrected Wilcoxon signed-rank tests, *p*_*corr*_ < 0.001). The latter indicates that the population of simultaneously-recorded units share information, at least to some degree. Thus, in response to natural sound streams, the auditory midbrain displays some degree of redundant information representation.

The degree of redundancy in unit pairs was quantified by computing the information interaction (*I*_*i*_) as the difference between the *I*_*sum*_ and the *I*_*joint*_ for each neuronal pair. *I*_*i*_ calculations can yield three possible outcomes: 1) redundant representations (*I*_*i*_ > 0, indicating shared information between units); 2) synergy (*I*_*i*_ < 0, both units provide bonus information when studied simultaneously); 3) independence (*I*_*i*_ = 0, units provide the same information considered both together and separated). Plotting *I*_*i*_ values vs. midbrain depth (mean depth of the two units forming each pair) revealed an exponential relation between these two variables, irrespectively of the stimulus presented to the bat (Fig. 5C). In other words, the highest *I*_*i*_ values (indicating more redundancy) are found in pairs of neurons recorded in deep midbrain layers. This trend was statistically validated by comparing redundancy across depth groups (Fig. 5D; FDR-corrected Wilcoxon rank-sum tests, p < 0.05). Taken together, our results suggest that the ventral IC provides more informative, but also more redundant representations of natural incoming communication and echolocation sound streams.

### Redundancy is highest in nearby units and arises from signal correlations in unit pairs

To unveil the origins of the redundant representations observed, we separated unit pairs according to whether they showed redundant or synergistic interactions (*I*_*i*_ > 0 and *I*_*i*_ <0, respectively). The *I*_*i*_ values were then analyzed considering the anatomical distance between the units forming the pairs (Fig. 6A-D). For this analysis, data from all stimulation sequences were pooled together. When considering only the redundant pairs, nearby units had higher redundancy levels than distant units (Fig. 6A). Even though the number of pairs decreased with inter-unit distance (see inset), statistical comparisons between nearby and faraway units were significant (Fig. 6B). In the case of synergistic pairs (Fig. 6C-D), we did not observe a clear dependence of *I*_*i*_ with depth although there was a small increase in *I*_*i*_ values for pairs in which the units were located far away from each other. In other words, it appeared as if information redundancy was more likely to occur when units were close to each other, while synergy tended to be higher in pairs of distant units.

**Figure 6.**
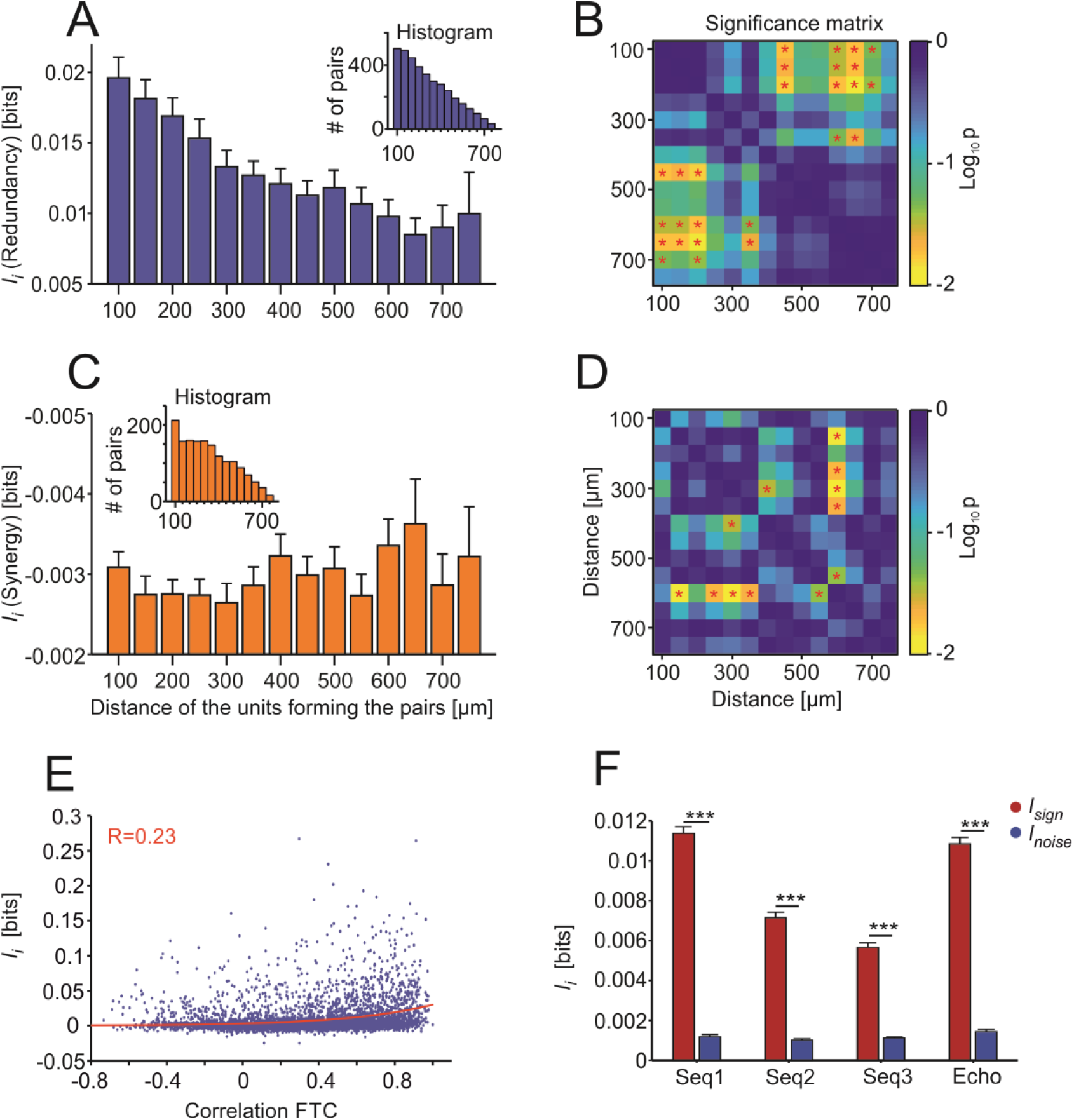
The redundancy is higher in nearby units and mostly comes from signal correlations. A) Bar plot of the redundancy values (mean + SEM) for those redundant pairs (i.e. that had *I*_*i*_ > 0) displayed according to the distance between the units that form the pairs. The inset shows the histogram of the pairs used for each distance. B) *P*-value matrix with a logarithmic scale. * *p*_*corr*_ < 0.05 for statistical comparisons from A. C) and D) with the same specifications for A and B, respectively, but for synergistic pairs (*I*_*i*_ < 0). E) Scatter plot of the *I*_*i*_ estimates shown against the correlation coefficients between the frequency tuning of the units forming the pairs. In red is the exponential curve fitted to the data with the corresponding correlation coefficient value. F) *I*_*i*_ broken down into signal (*I*_*sign*_; red) and noise (*I*_*noise*_; blue) correlations. Statistical comparisons performed by FDR-corrected Wilcoxon signed-rank tests. FTC: frequency tuning curve. *** *p*_*corr*_ < 0.001.

We also tested whether redundancy depended on units having similar iso-level frequency tuning curves. To that end, *I*_*i*_ was analyzed considering the Pearson correlation coefficients between the units forming the pairs (Fig. 6E). There was a moderate dependence between these two variables resulting in a correlation coefficient of 0.23 using an exponential fit (Fig. 6E). This shows a tendency towards higher correlated FTCs having more redundancy.

In a last step, we separated *I*_*i*_ into two components: 1) signal correlations (*I*_*sign*_), which represent the similarity between the average response in the two neurons studied across different time frames of the same stimulation sequence; and 2) noise correlations (*I*_*noise*_), which refer to the trial-by-trial variability in the responses (Averbeck et al., 2006). While *I*_*sign*_ always result in redundancy, *I*_*noise*_ can lead to either redundancy or synergy. Our data shows that in the IC, most of the redundancy between two units results from signal correlations (Fig. 6F). Thus, we can conclude that the redundancy observed in the IC is mostly stimulus-driven and does not necessarily represent an internal feature (noise) of the neuronal pairs.

## Discussion

In this study, we conducted simultaneous recordings of neuronal activity across the entire dorso-ventral extent of the inferior colliculus in awake bats presented with natural sound sequences. Our analysis focused on the spatial pattern of information representation at the midbrain level in response to natural sound streams. This is an important aspect for identifying which parts of the IC are instrumental for conveying information to other brain structures, such as the auditory thalamus, cortex and sensory-motor structures. Moreover, understanding how natural utterances are represented in the IC has direct translational implications, as this structure is a target area for prostheses aimed to help patients who cannot benefit from cochlear implants (Lim et al., 2009).

Our main findings are: 1) in bats, neurons carrying the most information about both distress and echolocation sequences are located ventrally in the IC; 2) unit pairs in ventral regions carry the highest redundancy as well; and 3) redundancy arises mostly from signal correlations in the units’ responses and is highest in nearby units with similar receptive fields.

### Ventral IC units have complex receptive fields and are highly informative about natural sound streams

In agreement with previous studies, we observed that the bat auditory midbrain contains units with double-peaked FTCs (Beetz et al., 2017; Casseday and Covey, 1992; Holmstrom et al., 2007; Mittmann and Wenstrup, 1995). In *C. perspicillata*, these units fire strongly to both low frequency (10-30 kHz) and high frequency sounds with a response notch in between (see example tuning curves in Fig. 2A and Fig. 3) and they are more likely to be found in ventral IC areas.

The fact that high-frequency units in the IC also respond to low frequencies has been described before in studies in other bat species such as the mustached bat, *Pteronotus parnelli* (Macías et al., 2012; Mittmann and Wenstrup, 1995; Portfors and Wenstrup, 1999). It appears that in some bat species, the tonotopic representation in the IC differs from the classical view so that, superimposed on the canonical dorso-ventral, low- to high-frequency axis, there is responsivity to low-frequencies among the high-frequency region. In *P. parnelli*, multi-peaked frequency tuning is especially useful for integrating information about biosonar call and echoes in different frequency channels during target-distance calculations (Macías et al., 2012; Mittmann and Wenstrup, 1995; O’Neill and Suga, 1982; Portfors and Wenstrup, 1999; Suga and O’Neill, 1979; Wenstrup et al., 1999). However, *C. perspicillata* (the species studied here) does not use multiple frequency channels for target distance calculations (Hagemann et al., 2010; Hechavarría et al., 2013; Kössl et al., 2014). Consequently, in this species, multi-peaked frequency tuning could offer advantages for representing communication sounds (i.e. distress) in widespread neuronal populations (Kanwal et al., 1994). Note that our data suggest that ventral IC neurons provide the highest information about distress and echolocation sequences. However, the latter does necessarily imply that dorsal IC areas do not respond to some of the features in the natural sounds.

According to our data, in the IC, neuronal information about natural acoustic sequences increase in an exponential continuum along the dorso-ventral axis. Ventral units convey the most information about both echolocation and communication calls. This was an unexpected result due to the spectral differences between echolocation and communication calls and the tonotopic organization of the IC. We argue that tracking natural sound streams could be related to the complexity of neuronal receptive fields. That is, the fact that ventral neurons have more complex receptive fields makes them more suited for providing information about the time course of natural sounds. According to this idea, in multi-peaked units, distress calls could activate inputs that correspond to both the low and high frequency peaks in the tuning curves.

Complex receptive fields could be beneficial for natural sound tracking because of several reasons. First, in response to distress, the simultaneous arrival of low- and high-frequency driven excitatory inputs would lead to spatiotemporal summation, which ultimately transduces into stronger responses (Magee, 2000). Another possibility that could be considered is that in response to distress sounds, adaptation in the low and high-frequency synapsis occurs in an asynchronous manner. In the latter scenario, ventral IC neurons would always receive excitatory inputs since adaptation alternates between low- and high-frequency information channels. Note, however, that the data gathered using echolocation sequences does not differ much from that gathered using distress sequences (Fig. 5). Echolocation does not carry strong energy at frequencies below 45 kHz and the latter suggest that high frequency inputs are sufficient for driving highly informative responses in the ventral IC. One could argue that echolocation requires specializations for precise temporal processing (Kössl et al., 2014; Neuweiler, 1990; Wenstrup and Portfors, 2011) and bats may profit from these adaptations even when listening to communication sounds. From the predictive coding framework (Ayala et al., 2016; Bastos et al., 2012; Remez et al., 1981), one could argue that echolocation processing implies a low-weighted prediction error which follows a “prior” hardwired in the system. In *C. persicillata* such neural prior occurs in the form of a good sound tracking ability in ventral IC areas. Communication-sound processing would benefit also from this innate high informative prior.

Note that our data offers only insights into the final activity output of IC units, but is not suited for assessing which of the above explanations (if any) contributes to the improved information representation in ventral IC layers. Information estimates used here only quantify the abilities of neurons to encode acoustic inputs, yet they do not capture the parameters of the stimulus the neurons are sensitive to (Borst and Theunissen, 1999; Chechik et al., 2006; Timme and Lapish, 2018). Thus, although both types of stimuli (distress and echolocation) showed similar patterns of information representation throughout the IC, different parameters of the two call types could contribute in different extents to the information maps observed.

### Possible origins of redundant information representation in the ventral IC

We observed that neurons in the ventral IC have complex receptive fields and carry high information content about natural sound sequences. However, the ventral IC also provides the most redundant information representations between units studied simultaneously. We show that redundancy in the ventral IC is linked to signal correlations, i.e. stimulus-induced activity correlations that arise when receptive fields overlap at least partially (Averbeck et al., 2006; Latham and Nirenberg, 2005). Common synaptic inputs can introduce both signal and noise correlations and could be the origin for the redundancy values reported here.

Our data indicates that in the IC, signal correlations are stronger than noise correlations, but this does not imply the absence of the latter. Signal correlations could relate to shared feedforward projections that dominate spiking during stimulus driven activity and to both the crossed projections from the contralateral IC and local connections. On the other hand, noise correlations can arise from common input as well, in combination with stimulus-independent neuromodulation acting on each neuron individually (Belitski et al., 2010). Noise correlations could also reflect feedback projections, e.g. from the auditory cortex (Jen et al., 1998; Yan and Suga, 1998) or the amygdala (Marsh et al., 2002), and they could regulate the IC’s processing at the single neuron level. Additionally, the IC receives crossed projections from the contralateral IC and has a dense network of intrinsic connections (Malmierca et al., 1995) that could also influence the information interactions.

In the present study, we report high redundancy levels in pairs formed by close-by neurons (~< 400 μm apart). This fact can be explained by the common inputs to neighbor neurons. Such common inputs might result from the IC’s tonotopy and they minimize wiring costs, an evolutionary adaptation linked to the formation of topographic maps (Chklovskii and Koulakov, 2001, 2004). In the bat IC, there are also “synergistic neuronal pairs” although consistent with previous literature (Narayanan et al., 2005; Samonds et al., 2004), the predominant form of information interaction is redundancy. Studies have argued that the main advantage of redundant information regimes is that multiple copies of essentially the same information exist in the neural network activity patterns, i.e. similar information channels exist (Pitkow and Angelaki, 2017).

The latter gives room to the implementation of computationally different transformations on each information channel. Such transformations might be used by the bat auditory system to extract relevant stream features that go beyond the representation of the sounds’ envelope (e.g. occurrence of bouts in distress sequences or precise coding of echo-delays (Beetz et al., 2016a; García-Rosales et al., 2018)) in upstream structures of the auditory hierarchy.

## Acknowledgements

The German Research Foundation (DFG) for funding this work and Gisa Prange for help with histological staining

## Author contributions

Conceptualization, J.C.H., F.G.R., L.L.J. and E.G.P.; Methodology, F.G.R. and J.C.H.; Investigation, E.G.P.; Formal Analysis: E.G.P. and F.G.R.; Writing – Original Draft, E.G.P. and J.C.H.; Writing-Review & Editing, J.C.H., F.G.R. and L.L.J.; Funding Acquisition, J.C.H; Supervision, J.C.H. and F.G.R.

## Declaration of Interests

The authors declare no competing financial interests

## Abbreviations

FTC: frequency tuning curve
IC: inferior colliculus
*I*_*rate*_: information of the spiking rate of a unit
*I*_*joint*_: *I*_*rate*_ calculated for a pair of units
*I*_*sum*_: sum of *I*_*rate*_ of the units forming a pair
*I*_*i*_: information interaction (*I*_*sum*_ − *I*_*joint*_ = *I*_*sign*_ + *I*_*noise*_)
*I*_*sign*_: signal correlations
*I*_*noise*_: noise correlations
SPL: sound pressure level

## STAR Methods

### Animals

For this study, 4 adult animals (3 males, species *C. perspicillata*) were used. The animals were taken from the bat colony at the Institute for cell biology and neuroscience at the Goethe University in Frankfurt am Main, Germany. The experiments comply with all current German laws on animal experimentation. All experimental protocols were approved by the Regierungspräsidium Darmstadt, permit #FU-1126.

### Surgical procedures

On the day of the surgery, the bats were caught at the colony and were anesthetized subcutaneously with a mixture of ketamine (10 mg/kg Ketavet, Pharmacia GmbH, Germany) and xylazine (38 mg/kg Rompun, Bayer Vital GmbH, Germany). Local anesthesia (ropivacaine hydrochloride, 2 mg/ml, Fresenius Kabi, Germany) was applied subcutaneously on the skin covering the skull. Under deep anesthesia, the skin in the dorsal part of the head was cut and removed, together with the muscle tissue that covers the dorsal and temporal regions of the skull. For fixation of the bat’s head during neurophysiology measurements, a custom-made metal rod (1-cm long, 0.1 cm diameter) was glued onto the skull using acrylic glue (Heraeus Kulzer GmbH), super glue (UHU) and dental cement (Paladur, Heraeus Kulzer GmbH, Germany). A craniotomy was performed 2-3 mm lateral from the midline above the lambdoid suture on the left hemisphere using a scalpel blade. The brain surface exposed was ~1 mm^2^.

During the surgery and the recordings, the custom-made bat holder was kept at 28° C with the aid of a heating pad. The surgery was performed on day 0, and the first recording was (at least) on day 2. Further recordings were performed on non-consecutive days. On each experimental day, experiments did not last longer than four hours, and during the recordings the animals received water every ~1.5 hours. The animals participated in the experiments for a maximum of 14 days. After this time period they were euthanized with an anesthetic overdose (0.1 ml pentobarbital, 160 mg/ml, Narcoren, Boehringer Ingelheim Vetmedica GmbH, Germany).

### Electrophysiological recordings

All recordings were performed in an electrically shielded and sound-proofed Faraday cage. Each recording consisted of three protocols: iso-level frequency tuning, spontaneous activity measurements and natural distress calls. In each recording day, the bat was placed on the holder and the rod on its skull was fixated to avoid head movements. Ropivacain (2 mg/ml, Fresenius Kabi, Germany) was applied topically whenever wounds were handled.

On the first recording day, a small hole in the skull was made for the reference and ground electrodes on the right hemisphere in a non-auditory area. The same electrode was used for these two purposes by short-circuiting their connectors. The recording electrode (A16, NeuroNexus, Ann Arbor, MI), was an iridium laminar probe containing 16 channels arranged vertically with 50 μm inter-channel distance, 1.1–1.4 MΩ impendence, 15 μm thickness, and a 50-μm space between the tip and the first channel. The electrode was introduced 2-3 mm laterally from the scalp midline, ~ 1 mm caudal to the lambdoid suture (Beetz et al., 2017; Coleman and Clerici, 1987), and perpendicularly to the surface of the brain, with the aid of a Piezo manipulator (PM- 101, Science products GmbH, Hofheim, Germany). Before starting the recordings, the electrode was lowered down by 1.1- 2.8 mm (depth measured from electrode’s tip). The tip’s depth was used as a reference to calculate the depth of all the channels. The position of the inferior colliculus was assessed by examining the responsivity to sounds across all recording channels. The sound used for testing for acoustic responsiveness was a short broadband distress syllable covering frequencies between 10-80 kHz. The electrophysiological signals obtained were amplified (USB-ME16-FAI-System, Multi Channel Systems MCS GmbH, Germany) and stored in a computer using a sampling frequency of 25 kHz. The data were stored and monitored on-line in MC-Rack (version 4.6.2, Multi Channel Systems MCS GmbH, Germany).

### Acoustic stimulation

For the present study three types of acoustic stimuli were used: pure tones, natural echolocation calls and distress calls from conspecifics. To assess the tonotopic arrangement of the IC, pure tones (10 ms duration, 0.5 ms rise/fall time) were presented at frequencies from 10 to 90 kHz in steps of 5 kHz at a fixed level of 60 dB SPL. The 17 pure tone stimuli were played in a pseudo-random manner with a total of 20 repetitions per sound.

The natural sounds comprised three distress and one biosonar sequences. The distress calls used as stimuli were recorded from conspecifics in the context of a previous study (see (Hechavarría et al., 2016a) for description of the procedures), as well as the echolocation call (Beetz et al., 2016b). These stimuli (also referred in this manuscript as *seq1*, *seq2*, *seq3* and *Echo*) had durations of 1.51, 2.47, 2.93 and 1.38 seconds, respectively. Each stimulus was played 50 times in a pseudo-random order. The root-mean-square level of the syllables that formed the sequences spanned between 74.5- and 93.1-dB SPL (Table 1). The sequences were multiplied at the beginning and end by a linear fading window (10 ms length) to avoid acoustic artifacts. Sounds were played from a sound card (ADI-2-Pro, RME, Germany) at a sampling rate of 192 kHz, connected to a power amplifier (Rotel RA-12 Integrated Amplifier, Japan) and to a speaker (NeoX 1.0 True Ribbon Tweeter; Fountek Electronics, China) placed 30 cm away from the right ear.

### Spike detection and sorting

Spikes were identified after filtering the data using a 3^rd^-order Butterworth band-pass filter with cutoff frequencies of 300 Hz and 3 kHz. The threshold for spike detection was 6 MAD (median absolute deviation). Spikes were sorted using the open-source algorithm SpyKING CIRCUS (Marre et al., 2018), a method that relies on density-based clustering and template matching, and can assign spikes clusters to individual channels in electrode arrays without cluster overlap. For further analysis, the cluster with the largest number of spikes was used for each channel. This spike-sorting algorithm ensures that the same cluster is not considered in different channels. Spike-sorted responses are referred to as “units” throughout the manuscript.

### Information theoretic analyses

All the information theoretic analysis was performed using the Information Breakdown Toolbox (ibTB) (Magri et al., 2009). The capability of a neuron with a set of responses *R* to encode a set of stimuli *S* can be quantified using Shannon’s mutual information (*I*(*R*;*S*)) (Shannon, 1948) using the following equation:

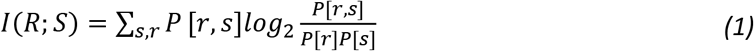

Where P[*s*] is the probability of presenting the stimulus *s*, *P*[*r*] is the probability of observing the spike count *r* and P[*r*,*s*] is the joint probability of presenting the stimulus *s* and observing the response *r*. The units of the mutual information are given in bits (when the base of the logarithm is 2). Each bit implies a reduction of the uncertainty about the stimulus by a factor of 2 by observing a single trial (Dayan and Abbott, 2001). One variable provides information about another variable when knowledge of the first, on average, reduces the uncertainty in the second (Cover and Thomas, 2006). Mutual information provides advantages in comparison to other methods as it is model independent and thus it is not necessary to hypothesize the type of interactions between the variables studied (Magri et al., 2009; Timme and Lapish, 2018) and captures all nonlinear dependencies in any statistical order (Kayser et al., 2009).

The naturalistic stimuli presented here were chunked into non-overlapping time windows, the neuronal responses to which were used to estimate the information, as it has been similarly done and described in other studies (Belitski et al., 2008; García-Rosales et al., 2018; Kayser et al., 2009; Montemurro et al., 2008; Steveninck et al., 1997). The time window considered here for the substimuli (T = 4 ms) has been selected to make our calculations comparable to those from studies in the AC (García-Rosales et al., 2018; Kayser et al., 2009). In order to calculate the information contained in the firing rate of each unit (*I*_*rate*_), the number of spikes that occurred in response (*r*) to each substimulus (*s*_*k*_) was determined. The responses were binarized, i.e. they show if there is at least one spike (*1*) or none (*0*), *r* = *{0, 1}*. *P*(*r*) indicates the probability of firing (or not) and was estimated considering all the 50 trials of each sequence.

Information was quantified by two main neuronal codes: the rate code (*I*_*rate*_) and the information carried by the rate of two units (*I*_*joint*_) recorded simultaneously. The *I*_*joint*_ was calculated in the same manner as the *I*_*rate*_ with the difference that now the response (*r*) can take four forms (instead of two) as it keeps track of which neuron fired, therefore *r = {0-0, 0-1, 1-0, 1-1}*. As mentioned in the preceding text, the spike-clustering algorithm used in this paper considers the geometry of the laminar probe. Therefore, for each recording channel, a different spike waveform is allocated. To further make sure that a unit was not paired with itself during *I*_*joint*_ calculations, we considered only pairs of units from channels separated by at least a 100-μm distance. The mutual information was calculated as well for groups of three, four and five units recorded simultaneously.

To evaluate if the information carried by a unit pair was independent, redundant or synergistic, we calculated the information interaction (*I*_*i*_). *I*_*i*_ in pairs of neurons can be quantified by the sum of information conveyed by those neurons individually (*I*_*sum*_ = *I*_*rate*(*a)*_ + *I*_*rate*(*b*)_) and the difference between information conveyed by the two neurons (*I*_*joint*_) (Brenner et al., 2000; Chechik et al., 2006; Narayanan et al., 2005):

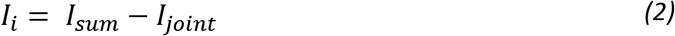

If *I*_*joint*_ < *I*_*sum*_ or simply if the *I*_*i*_ estimate is positive, the units carry redundant information; if *I*_*joint*_ = *I*_*sum*_, the units carry independent information and if *I*_*joint*_ > *I*_*sum*_, or if the *I*_*i*_ estimate is negative, they carry synergistic information. This compares the information available in the joint response to the information available in the individual responses.

The *I*_*i*_ was broken down into two components: the effects of signal and noise correlations. The signal similarity component (*I*_*sign*_) quantifies the amount of information specifically due to signal correlations, i.e. the degree to which the (trial-averaged) signal changes with the stimulus (Belitski et al., 2008). The noise correlation component (*I*_*noise*_) quantifies the trial-by-trial variability (Magri et al., 2009).

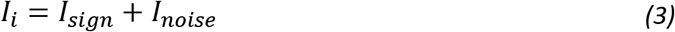

Information estimates were calculated by the “direct” method (Borst and Theunissen, 1999), which requires a large amount of experimental data as it does not make any assumption about response probability distributions. As it is very difficult and improbable to observe all possible responses from the entire response set (*R*) (Panzeri and Treves, 1996; Strong et al., 1998) due to the lack of unlimited number of trials, the quantities calculated with the estimated probabilities will always be biased. To account for that, the ibTB toolbox (Magri et al., 2009) uses the Quadratic Extrapolation (QE) procedure (Strong et al., 1998) and the subtraction of any remaining bias by a bootstrap procedure (Montemurro et al., 2008). In addition, for the *I*_*joint*_ tests, the Shuffling procedure (Panzeri et al., 2007) was also applied; which is also implemented in the ibTB toolbox.

In order to test the performance of the bias correction, simulated data with first order statistics close to those of the real data were generated. For the *I*_*rate*_, spike responses were generated (inhomogeneous Poisson processes) with the same PSTH as each real unit used for the analysis (as in García-Rosales et al., 2018). Information was computed for the simulated data using the same parameters than for the original data, for all the neural codes used (*I*_*rate*_, and *I*_*joint*_ for pairs, triplets, quadruplets and quintuplets) and for different number of trials (4, 8, 16, 32, 50, 64, 128, 256, 512). According to our results of the performance of the bias correction on simulated data, the bias was negligible for the *I*_*rate*_, *I*_*joint*_ for pairs and triplets and had slightly negative for the *I*_*joint*_ for quadruplets and quintuplets, the information estimates underestimate the true information values for the last two variables.

## Supplemental Information

**Figure S1.**
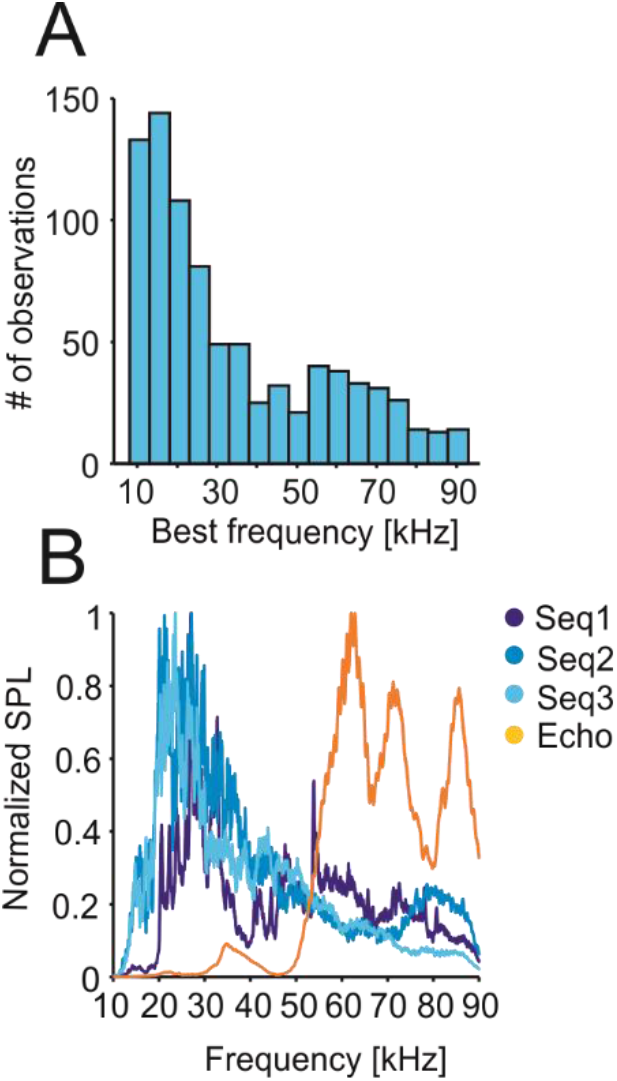
The best frequencies of most of IC’s neurons match the frequency range that has the most amplitude in distress calls. A) Histogram of the best frequencies for all the units used in further analyses (n = 864). B) Spectra of the natural calls used in this study with normalized SPL.

**Figure S2.**
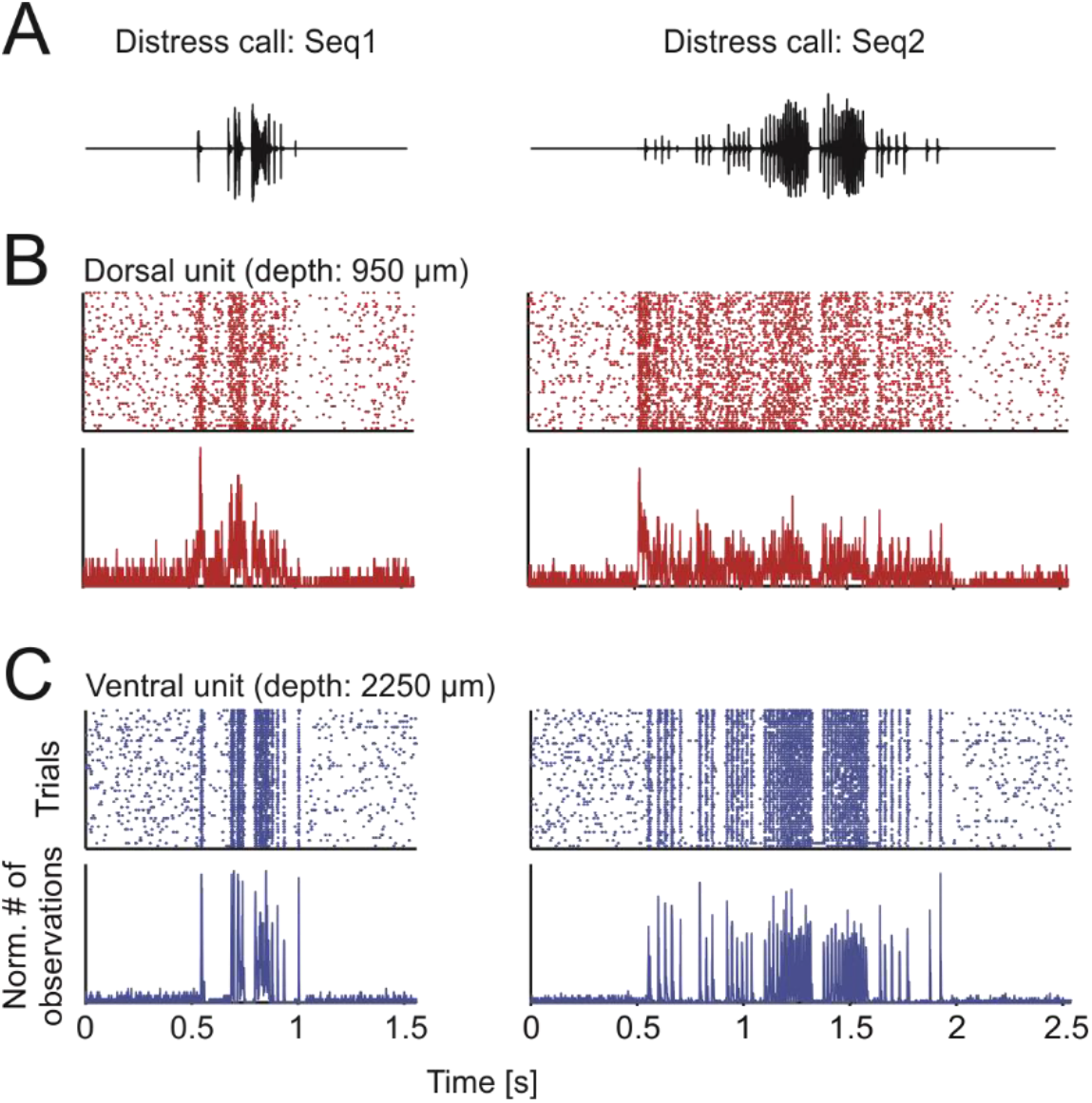
Neural activity of dorsal and ventral units for Seq1 and Seq2. A) Oscillograms of two of the natural calls used as stimuli. B) Raster plots (50 trials in total; top) and peristimulus time histogram (PSTH; 1 ms precision; bottom) of one exemplary dorsal unit in response to sequences shown in A. C) Same specifications as in B but for an exemplary ventral unit (both are the same as in Fig. 3. Note the higher precision and reliability in the ventral unit.

**Figure S3.**
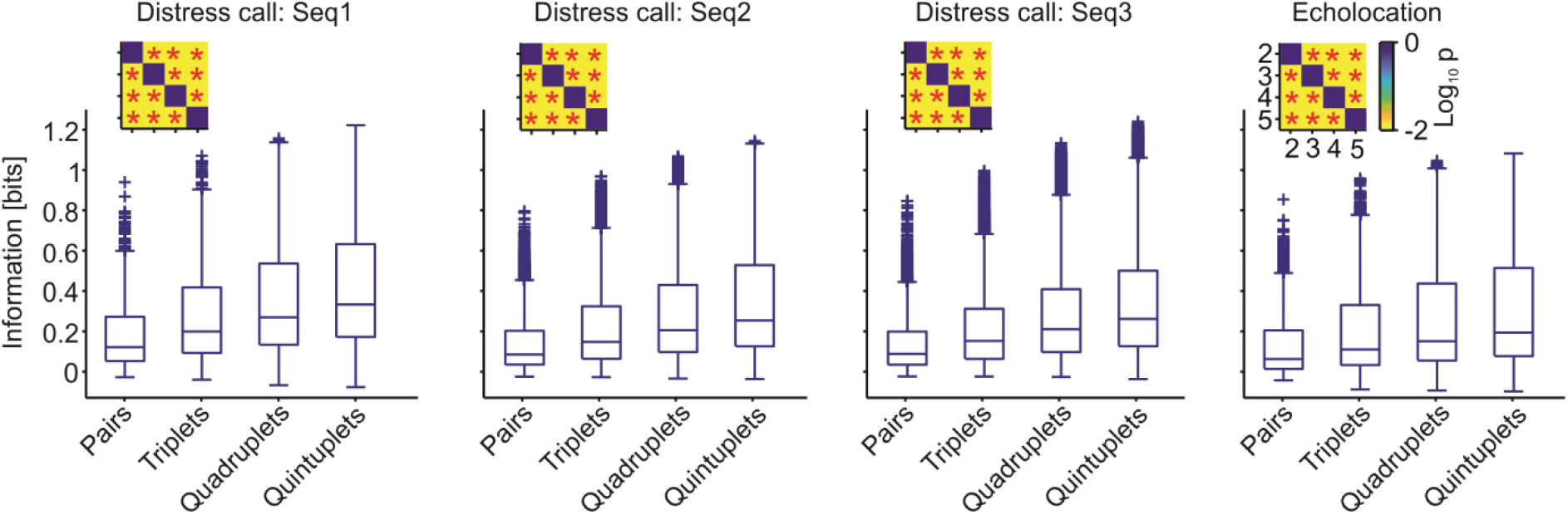
Mutual information for groups of units. Boxplots of the mutual information carried by groups of units (pairs, triplets, quadruplets and quintuplets). Note that only were considered units that were recorded from electrodes distances at least 100 μm.

